# Epigenomic divergence underlies sequence polymorphism and the evolutionary fate of duplicate paralogs in *A. thaliana*

**DOI:** 10.1101/2023.03.02.530917

**Authors:** Sunil K. Kenchanmane Raju, Mariele Lensink, Daniel J. Kliebenstein, Chad Niederhuth, Grey Monroe

## Abstract

Processes affecting rates of sequence polymorphism are fundamental to molecular evolution and the evolutionary fate of gene duplicates. The relationship between gene activity and sequence polymorphism can influence the likelihood that functionally redundant gene copies are co-maintained in stable evolutionary equilibria versus other outcomes such as neo-functionalization. Here we investigate genic variation in epigenome-associated polymorphism rates in *Arabidopsis thaliana* and consider whether these affect the evolution of gene duplicates. We compared the frequency of sequence polymorphism and patterns of genetic differentiation between genes classified by exon methylation patterns: unmethylated (unM), gene-body methylated (gbM), and transposon-like methylated (teM) states, which reflect divergence in gene expression. We found that the frequency of polymorphism was higher in teM (transcriptionally repressed, tissue-specific) genes and lower in gbM (active, constitutively expressed) genes. Comparisons of gene duplicates were largely consistent with genome-wide patterns - gene copies that exhibit teM tend to accumulate higher sequence polymorphism, evolve faster, and are in chromatin states associated with reduced DNA repair. This relationship between expression, the epigenome, and polymorphism may lead to the breakdown of equilibrium states that would otherwise maintain genetic redundancies. Epigenome-mediated polymorphism rate variation may therefore aid the pseudogenization of duplicate paralogs or increase the evolution of novel gene functions in duplicate paralogs maintained over evolutionary time.

## Introduction

DNA sequence polymorphism is the ultimate source of variation in evolution. The nature of sequence polymorphism between genes also plays a central role in the evolutionary dynamics of gene duplicates (Panchy et al. 2016). Following gene duplication, selection may be relaxed on one or both paralogs to allow mutation accumulation leading to neo-, sub-functionalization, or pseudogenization (Brookfield 1992; Panchy et al. 2016; Kuzmin et al. 2022). The maintenance of gene duplicates with redundant functions has also been described, with the extent and evolutionary conditions necessary to maintain truly redundant duplicates being a long-standing question (Pickett and Meeks-Wagner 1995; Wagner 2002; Kuzmin et al. 2022). Nowak et al. (1997) showed that divergence in sequence polymorphism between paralogous pairs could disrupt equilibria that maintain redundant functions, leading to an accelerated loss of duplicates if novel functions do not evolve. Besides cases where selection favors the increased dosage from multiple gene copies, maintenance of redundancy among gene pairs is expected to occur under three distinct models: 1) when sequence polymorphism (mutation) rates are equal between paralogs, 2) when sequence polymorphism and gene activity change together such that decreased activity of a paralog is accompanied by a reduction in its sequence polymorphism or 3) one paralog evolves an additional secondary function (neo-functionalization) (Ohno 1970; Nowak et al. 1997; Lynch and Conery 2000). Characterizing the forces that affect the rate of polymorphism, especially if they give rise to correlations between the frequency of polymorphism and activity, is thus critical for explaining the evolutionary fate of duplicated genes (Rodin and Riggs 2003).

Among the essential features for predicting the redundancy of gene pairs, gene expression profiles, gene function, and mode of gene duplication were more informative than evolutionary rate-related features, such as having a paralog from a recent duplication event (Cusack et al. 2021). Thus, gene expression variability during development and/or environmental challenges can influence the evolutionary trajectories of duplicate genes. Roberts and Josephs (2022) have shown that highly treatment-specific genes in *A. thaliana* belong to large gene families, suggesting that gene duplication and following evolutionary fates of duplicates may be associated with the evolution of expression specificity (Van de Peer et al. 2009; Panchy et al. 2016; Roberts and Josephs 2022)

The factors influencing the frequency of polymorphism between genes and whether the gene-level variation is neutral or adaptive are questions of recently renewed investigation. On the one hand, variation between genes may be the outcome of neutral stochastic processes, with differences arising irrespective of gene activity, function, or contribution to fitness. In this case, the frequency of polymorphism and gene activity are expected to change independently, providing opportunities for duplicate genes to be maintained by the equilibrium state under model 2 described by Nowak et al. (1997) (Nowak et al. 1997). Such equilibria states would be even more common if the sequence polymorphism were deleterious, for example, in the case that transcription itself is mutagenic since genes with higher mutation rates will accumulate polymorphism faster (Lynch et al. 2016).

On the other hand, variation in polymorphism rates could be adaptive, reflecting mechanisms that link gene activity and function with the rate of polymorphism in beneficial directions (i.e., if active genes experience reduced mutation rate, and vice-versa). Such non-randomness is unlikely to evolve on a gene-by-gene basis because the efficacy of selection at that scale is inadequate to overcome the barrier imposed by drift (Lynch 2010; Hodgkinson and Eyre-Walker 2011; Stoletzki and Eyre-Walker 2011; Lynch et al. 2016) However, the drift-barrier can be overcome if the rate of polymorphism is linked to the epigenome (Martincorena and Luscombe 2013), allowing the evolution of systemic bias in accumulation of polymorphism in relation to gene function and activity characterized by shared epigenome states. Mutation rates are not independent of the epigenome across diverse organisms (Schuster-Böckler and Lehner 2012; Li et al. 2013; Makova and Hardison 2015; Supek and Lehner 2017; Habig et al. 2021; Monroe et al. 2022; Quiroz et al. 2022), suggesting the potential for mutation modifier systems relevant for understanding the evolutionary fate of gene duplicates. Specifically, adaptive polymorphism rate variation is expected to reduce the chances of equilibrium states in which genes with lower activity also have lower polymorphism (fewer cases of model 2 described by Nowak et al. (1997)).

One potential epigenomic mark that could differentiate duplicated genes is the expression regulatory state that is highly variable between duplicated genes (Keller and Yi 2014; Wang et al. 2014; Kim et al. 2015; Raju et al. 2022). In addition, duplicate paralogs often evolve asymmetrically, with one paralog directed towards repressed expression (Casneuf et al. 2006; Ganko et al. 2007; Panchy et al. 2016; Gillard et al. 2021). However, whether such divergence in expression is random with respect to changes in sequence polymorphism remains unknown. (Keller and Yi 2014; Wang et al. 2014; Kim et al. 2015)

Epigenomic features reflect the expression-related regulatory states of genes (Feng and Jacobsen 2011; He et al. 2011; Kawakatsu et al. 2016; Agarwal et al. 2020; An et al. 2020; Zhao et al. 2020; Lloyd and Lister 2022). In plants, cytosine methylation occurs in three different nucleotide contexts: dinucleotide CG sites and trinucleotide CHG and CHH (H = A, T, or C) sites (Gruenbaum et al. 1981; Finnegan and Dennis 1993; Meyer et al. 1994) Differing patterns of methylation in these contexts distinguish genes with constitutive versus silent or tissue-specific expression patterns, thus marking genes maintained in different and predictable epigenomic states (Niederhuth and Schmitz 2017).

Genes whose exons are enriched for CG methylation only (known as gene body methylation or **gbM genes**) tend to be constitutively expressed and are associated with other epigenomic features such as H3K4me1 that are characteristic of active genes (Zhang et al. 2009; Takuno and Gaut 2013; Bewick et al. 2016). In contrast, genes with both CG and non-CG (CHH and CHG) methylation (known as transposon-like methylation or **teM genes**) tend to be silent or expressed only in specific tissues and are correlated with epigenomic features associated with transcriptional silencing like H3K9me2 (Kawakatsu et al. 2016; Niederhuth et al. 2016; Inagaki et al. 2017). Finally, genes lacking exonic methylation (known as unmethylated or **unM genes**) tend to exhibit patterns of expression and epigenome features that reflect neither broad activation nor silencing. Comparison of genes in these alternative classes of genic methylation provides an opportunity to test hypotheses about the rates of polymorphism and their effects on the divergence of gene expression regulatory status.

Here we set out to ask if the frequency of polymorphism differs with respect to degrees of gene activation, defined according to the distinct epigenomic states. We first examined patterns at genome-wide scales and later focused specific analyses on duplicate paralogs. We find that active genes show lower polymorphism than silenced genes, indicative of joint influences of mutation rate variation and selection, a pattern with implications for the evolutionary fate of duplicated genes.

## Materials and Methods

### Genic methylation classification

Genome and gene annotations of the *Arabidopsis thaliana Col-0* accession used in this study were derived from Araport11 (Cheng et al. 2017). We implemented a custom filtering step to remove potentially mis-annotated transposons as described before (Bowman et al. 2017) with necessary modifications. First, using ‘*hmmscan*’ from the HMMER software package (Potter et al. 2018), each query gene was scanned against Pfam A HMM profile filtering genes matching a curated list of TE domains with an e-value <1e-5 (https://github.com/Childs-Lab/GC_specific_MAKER). We then performed a *DIAMOND blastP* (Buchfink, Xie, and Huson 2015a) against the transposase database (www.hrt.msu.edu/uploads/535/78637/Tpases020812.gz), removing hits with an e-value < 1e-10. This removed a total of 357 potentially misannotated genes. Previously published whole-genome bisulfite sequencing (WGBS) dataset (SRX2511763, (Picard and Gehring 2017)) was mapped to the *A. thaliana* genome using *methylpy v1.2.9 (Schultz et al*. *2015)*. For each gene, a binomial test was performed to test for enrichment of methylation in CG, CHG, and CHH contexts (H = A, C, or T) in the coding region of primary transcripts against background methylation rates calculated for all protein-coding genes across 43 plant species (Niederhuth et al. 2016; Raju et al. 2022). Benjamini-Hochberg’s false discovery rate (FDR) correction was applied to all p-values (Benjamini and Hochberg 1995). Genes were classified into three categories based on their genic methylation as previously described (Takuno and Gaut 2012; Niederhuth et al. 2016; Raju et al. 2022). Genes were classified as unmethylated (unM) if they had ≤ 1 methylated site in any context or ≤ 2% weighted methylation (Schultz et al. 2012) for all contexts (CG, CHG, and CHH). Genes with enrichment for CG methylation with ≥ 10 CG sites and no significant enrichment for CHG and CHH methylation were classified as gene-body methylated (gbM) genes. Genes with ≥ 10 sites in any of the three contexts and HG and CHH methylation enrichment were classified as transposon-like methylated (teM). Genes with intermediate DNA methylation levels that could not be confidently categorized into the above three categories were considered ‘unclassified.’ Genes lacking sufficient DNA methylation data were considered ‘missing’ and denoted as ‘NA’. Previously processed cytosine methylation files from WGBS data of 928 accessions from the 1001 *A. thaliana* epigenomes project were downloaded (GSE43857, (Kawakatsu et al. 2016)). All genes in each accession were classified as described above, and the frequency of unM, gbM, and teM epiallele for each gene in the population was calculated using a custom R script.

### Genetic diversity in a global set of *A. thaliana* accessions

Rates of single nucleotide polymorphism (SNPs) and insertion/deletion (indel) polymorphism were calculated for exons in TAIR10 gene models from sequence polymorphism dataset across 1,135 natural accessions of *A. thaliana (Alonso-Blanco 2016)*. For each gene, the percentage of nucleotides with polymorphism observed in the natural populations within 100bp windows spanning the gene was determined as previously described (Monroe et al. 2022). A one-way ANOVA test was performed to test for differences between genic methylation classifications and rates of polymorphism across the gene. A Spearman rank-order correlation or Spearman’s rho (⍴) was used to determine the relationship between the rates of polymorphism and epiallele frequency within the population.

### Test of neutrality using population genetic test statistic Tajima’s D

A site frequency spectrum metric, Tajima’s D, was analyzed for each gene to quantify the relative importance of mutation rates and selection in shaping the rates of sequence evolution. Tajima’s statistic is computed as a measure of the mean number of pairwise differences and the total number of segregating sites (Tajima 1989). Tajima’s D was calculated on 100 bp windows across the genome, and values were averaged for each gene based on their TSS and TTS. The confidence interval (±2 SEM) around the mean value was calculated using bootstrapping (n=100) as previously described (Monroe et al. 2022). Theory predicts that the mutation rate is negatively correlated with Tajima’s D but positively correlated with polymorphism rates. In contrast, the strength of purifying selection is negatively correlated with both Tajima’s D and polymorphism rates. A more negative Tajima’s D value suggests sequences with a higher fraction of sites under purifying selection and enrichment of rare alleles. Thus a joint analysis of Tajima’s D and rates of polymorphism can provide insights into the underlying drivers influencing rates of sequence evolution.

### Simulations of Tajima’s D

To contextualize empirical observations of epigenome-associated polymorphism divergence we simulated a 3,000 bp sequence under different combinations of multipliers of the standard baseline mutation rate (1e^-06^ mutations per base pair per generation (Ossowski et al. 2010) and different proportions of deleterious mutations. We ran each simulation on a population of 1,000 individuals for 10,000 generations in replicates of 300. Three different scaled mutation rates of 0.2, 1, and 1.5 reflecting an 80% reduction, 0%, and 50% increase in mutation rates relative to the baseline, were used. Similarly, proportions of deleterious mutations were assigned to 0, 0.1, 0.3, and 0.5, reflecting 0%, 10%, 30%, and 50% deleterious mutations. To determine the consequences of altering the distribution of fitness effects (DFE) of each mutation, additional simulations were performed using the same experimental setup, with changes to the scaled mutation rates (50% and 0% reduction) and proportions of deleterious mutations (10%, 30%, and 50%). The DFE of each deleterious mutation was represented by a gamma distribution, with a new simulation for each of the following mean values of the distribution: -0.5,-0.3,-0.1,-0.05,-0.01,-0.005,-0.001, and 0. In addition, other parameters, including recombination and selfing rates, were determined empirically from the literature (Bomblies et al. 2010; Platt et al. 2010). Average values for Tajima’s D and polymorphism were then calculated across the 300 replicates for each unique parameter combination.

### Identification of duplicate paralogs with divergent methylation

Protein-coding genes in *A. thaliana* were classified into different types of gene duplicates based on whether they were derived from whole-genome duplication or one of the four types of single gene duplicates (tandem, proximal, translocated, and dispersed). DIAMOND was used to perform ‘blastp’ for the target genome, *A. thaliana*, with itself and the outgroup genome *Amborella trichopoda* (Amborella *Genome* Project 2013; Buchfink *et al*. *2015*), retaining blast hits with e-value < 1e^-5^. *Duplicate gene classification was performed by DupGen_finder_unique* pipeline with the options -s 5 (requiring ≥ 5 genes to call a collinear block) and -d 10 (≤ 10 intervening genes to call ‘proximal’ duplicates). BLAST hits were filtered to remove hits from different orthogroups. Within gene families, pairs with the lowest e-value were retained to avoid over-counting. Finally, duplicate gene pairs for each of the five types of gene duplications were merged with genic methylation classifications of *A. thaliana* genes. Only paralogs with divergent methylation were used in the downstream analysis.

### Percent polymorphism differences between paralogs with divergent methylation

For duplicate paralogs with divergent methylation, the percentage of nucleotides with polymorphism observed in the natural populations within 100bp windows spanning the gene was determined for paralogs 1 and 2. A t-test was performed for differences in means of percent polymorphism and Tajima’s D values in duplicate pairs with methylation divergence (unM-gbM, unM-teM, and gbM-teM pairs). Unfortunately, we can not determine the parental gene and the newly duplicated copy for all these duplicate pairs. Thus the directionality of methylation divergence and polymorphism changes cannot be ascertained. However, in the case of translocated duplicates, we can ascertain the parental gene and the newly derived translocated copy using synteny information (Wang et al. 2012). Genic methylation and frequency of polymorphism in parental and translocated copies were obtained using custom R scripts.

### PDS5C and H3K4me1 ChIP-seq data

ChIP-seq datasets for H3K4me1 in *Arabidopsis thaliana* (Col-0 genetic background) were obtained from the Plant Chromatin State Database (Liu et al. 2018). H3K4me1 levels were calculated for each gene exon, with individual datasets being scaled to a mean of 0. Values were averaged across datasets for each gene for the final H3K4me1 enrichment score. PDS5C ChIP-seq data were previously published (Niu et al. 2021). Enrichment was calculated as previously described by Niu et al, (2021) from two replicates, which were averaged across the gene body of each gene as log2[(1 + n_ChIP)/N_ChIP] – log2[(1 + n_Input)/N_Input)], where n_ChIP and n_Input represent the total depth of mapped ChIP and input fragments in a region, and N_ChIP and N_Input are the numbers of total depths of unique mapped fragments.

## Results

### Identifying epigneomically distinct genes

We classified all protein-coding genes in *Arabidopsis thaliana Col-0* into three categories based on their genic methylation status as described previously (**Figure 1a**) (Raju et al. 2022). Genes enriched for CG-only methylation were classified as gene-body methylated (gbM), while genes with CG and non-CG methylation were classified as transposon-like methylated genes (teM). Genes without DNA methylation in any context were classified as unmethylated (unM). Of the 27,444 protein-coding genes, 357 genes were removed based on our stringent custom filtering step (see methods). Among the remaining 27087 genes, more than half, 15,647, were classified as unM, while 4,128 genes were classified as gbM and 1,289 genes as teM (**Table S1**). We checked the expression profiles of genes classified as teM to confirm that they are actual protein-coding genes and not mis-annotated transposons or pseudogenes. More than 98.4% of teM genes (1269 genes) showed mRNA expression in at least one tissue/treatment in the *A. thaliana* gene expression atlas (Klepikova et al., 2016). The set of 27,087 genes analyzed does not contain any genes annotated as transposable-element genes and/or pseudogenes, indicating that teM genes are mostly protein-coding genes with detectable expression in at least a few tissues/treatments. H3K4me1, which highly correlated with CG methylation in transcribed genic regions(Zhang et al. 2009), showed higher density in gbM genes, while teM genes had the lowest H3K4me1 density. GbM and unM genes had a lower density of H3K9me1 and H3K9me2 than teM genes (**Figure S1**), consistent with their transcription-coupled replacement in actively transcribed genes(Johnson et al. 2002). These results suggest that the distinct DNA methylation states (unM, gbM, and teM) distinguish genes differentiated by broader epigenomic states, including histone modifications.

**Figure 1:**
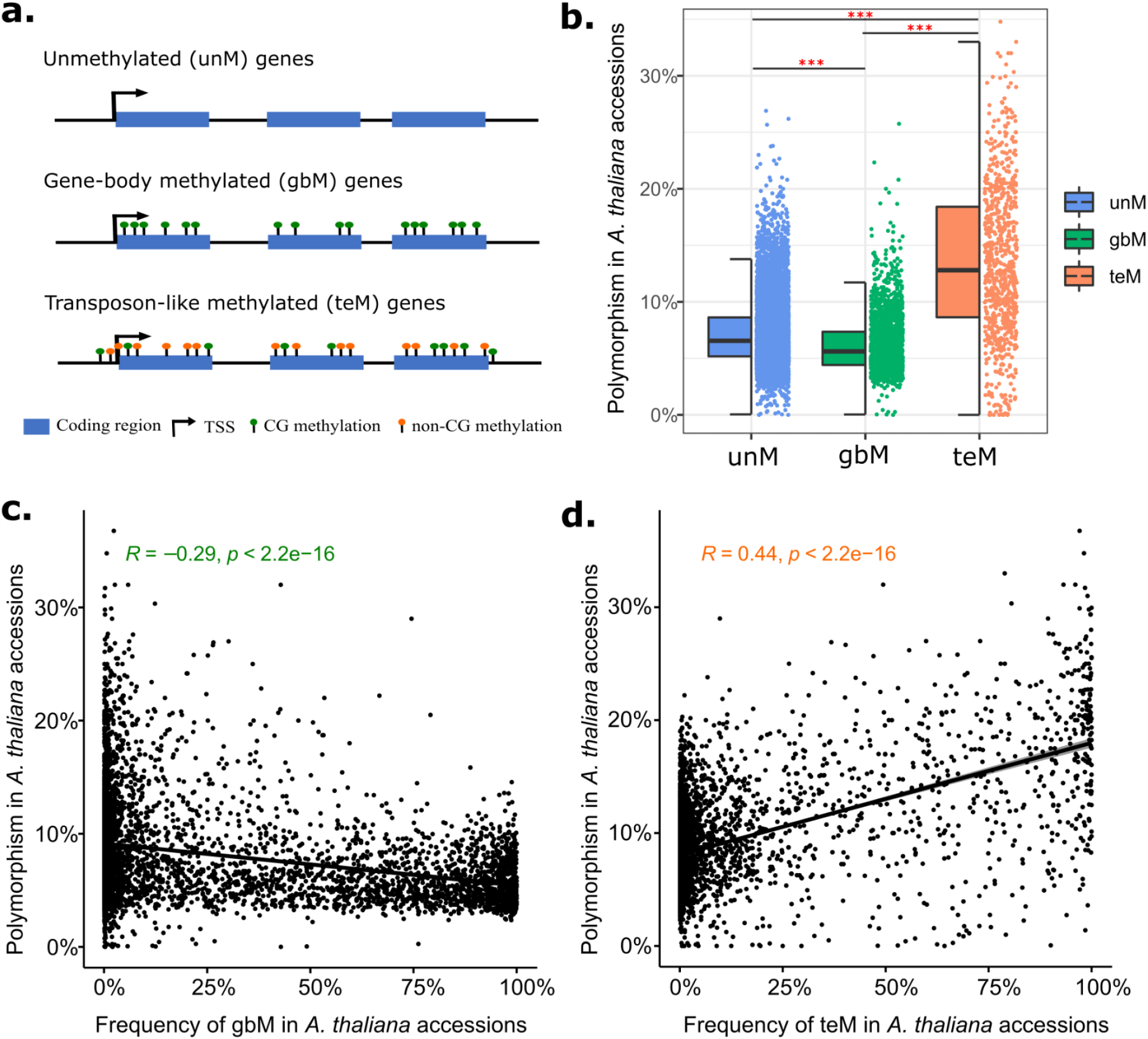
Genie methylation and its association with single-nucleotide polymorphisms in *A. thaliana* population. Schematic representation of the three different genie methylation classification (unM-unmethylated, gbM - gene-body methylated, and teM - transposon-like methylated). Green pins represent CG methylation while the orange pins represent non-CG methylation (mCHG and mCHH) on coding regions (blue rectangles) of genes. B) Percentage of single-nucleotide polymorphisms across *A. thaliana* accessions and their genie methylation classification in the *A. thaliana* accession *Co/-0*. *** represent statistically significant difference in SNP percentages among unM, gbM, and teM genes following a one-way ANOVA test at p-value < 0.001 C) and D) Spearman’s rank order correlation between epiallele frequency (gbM and teM) and single-nucleotide polymorphismsin the natural accessions of *A. thalina*. Negative Sprerman’s rho (r_s_) is indicated in green (C), while positive correlation is depicted in orange (D).

### Association between epigenomic states and rates of sequence evolution

To test whether the DNA methylation patterns of genes influence the differential accumulation of sequence polymorphism in natural populations, we looked at polymorphisms across natural accessions of *A. thaliana*. Polymorphism among natural *A. thaliana* accessions was extracted from the 1001 Genomes collection (Alonso-Blanco 2016), and the percentage of nucleotides with polymorphism within 100 bp windows was calculated for each gene. We found significant differences in polymorphism between the three genic methylation classes (One-way ANOVA, f(2)=3556, p-value < 0.001; **Figure 1b**). GbM genes and teM genes showed the lowest and highest rates of polymorphism, respectively, while unmethylated genes showed intermediate rates of polymorphism. TeM genes showed 1.37X (8.5%) and 1.06X (7.6%) more polymorphism than gbM and unM genes (Tukey’s test, adj p-value < 0.001). UnM genes showed 0.15X (0.9%) higher polymorphism in the natural population compared to gbM genes (Tukey’s test, adj p-value < 0.001).

Because methylation states can differ between accessions, we wanted to check that the patterns observed are not unique to epigenome states in Col-0. We examined methylomes of 928 *A. thaliana* accessions from the 1001 Epigenomes Project (Kawakatsu et al. 2016) and calculated the frequency of gbM/unM/teM alleles in the population. This enabled us to determine the relationship between the frequency of epiallele and SNPs in the natural *A. thaliana* population. The frequency of gbM within the population showed a weak negative correlation with SNPs (r_S_ = -0.29, p-value < 0.001, **Figure 1c**). In contrast, teM frequency in the population showed a moderate positive correlation with SNPs (r_S_ = 0.44, p-value < 0.001, **Figure 1d**). We also saw a weak but significant negative correlation between SNP polymorphism and unM frequency (r_S_ = -0.076, p-value < 0.001, **Figure S2**). Genes with a higher frequency of teM across the population tend to accumulate more polymorphism. In comparison, genes with a higher frequency of gbM show a lower proportion of polymorphism across the population. These results suggest that distinct epigenomic features in unM, gbM, and teM genes are associated with the accumulation of polymorphism in genes at the population level.

### Characterizing selection and mutation rate in natural populations in relation to epigenome state

Rates of sequence variability in natural populations are influenced by natural selection. Hence, the accumulation of polymorphism alone is insufficient to infer the evolutionary forces acting on genes with different methylation states. We calculated summary statistics for signatures of selection for each *A. thaliana* gene using their synonymous (Ds) and non-synonymous (Dn) substitution divergence from *Arabidopsis lyrata*. Natural sequence evolution in gbM genes showed lower Dn/Ds ratios compared to unM and teM genes, consistent with the expected stronger purifying selection on genes with housekeeping functions. On the other hand, genes classified as teM genes showed higher Dn/Ds ratios compared to unM and gbM genes suggesting that these genes are accumulating more non-synonymous mutations and could be under more relaxed purifying, fluctuating, positive, or diversifying selection (**Figure S3**).

Toward determining the relative effects of mutation rate versus selection on rates of sequence evolution (levels of polymorphism), we analyzed Tajima’s D, which is affected by selection and mutation rate (Tajima 1989). Tajima’s D tends to be more negative as a function of the number of selected sites under purifying selection (proportion of sites). Thus, our observation that gbM genes have more negative Tajima’s D than unM genes (**Figure 2a**) suggests that they experience more sites under purifying selection, which is expected since the gbM state reflects constitutive housekeeping expression states, consistent with their lower Dn/Ds ratio. In natural populations where the proportion of neutral sites is <100%, more negative Tajima’s D can also be caused by elevated mutation rates (Morales-Arce et al. 2020). Our analysis showed that Tajima’s D is more negative in teM genes compared to unM and gbM genes (One-way ANOVA, f(2)=235.8, p-value < 0.001). However, it is unlikely that this is caused by teM genes having more sites under purifying selection, as they have a higher level of polymorphism than gbM and greater Dn/Ds (**Figure S3**). Instead, the greater polymorphism combined with a more negative Tajima’s D value is consistent with the expected effects of elevated mutation rates and a weaker DFE.

**Figure 2:**
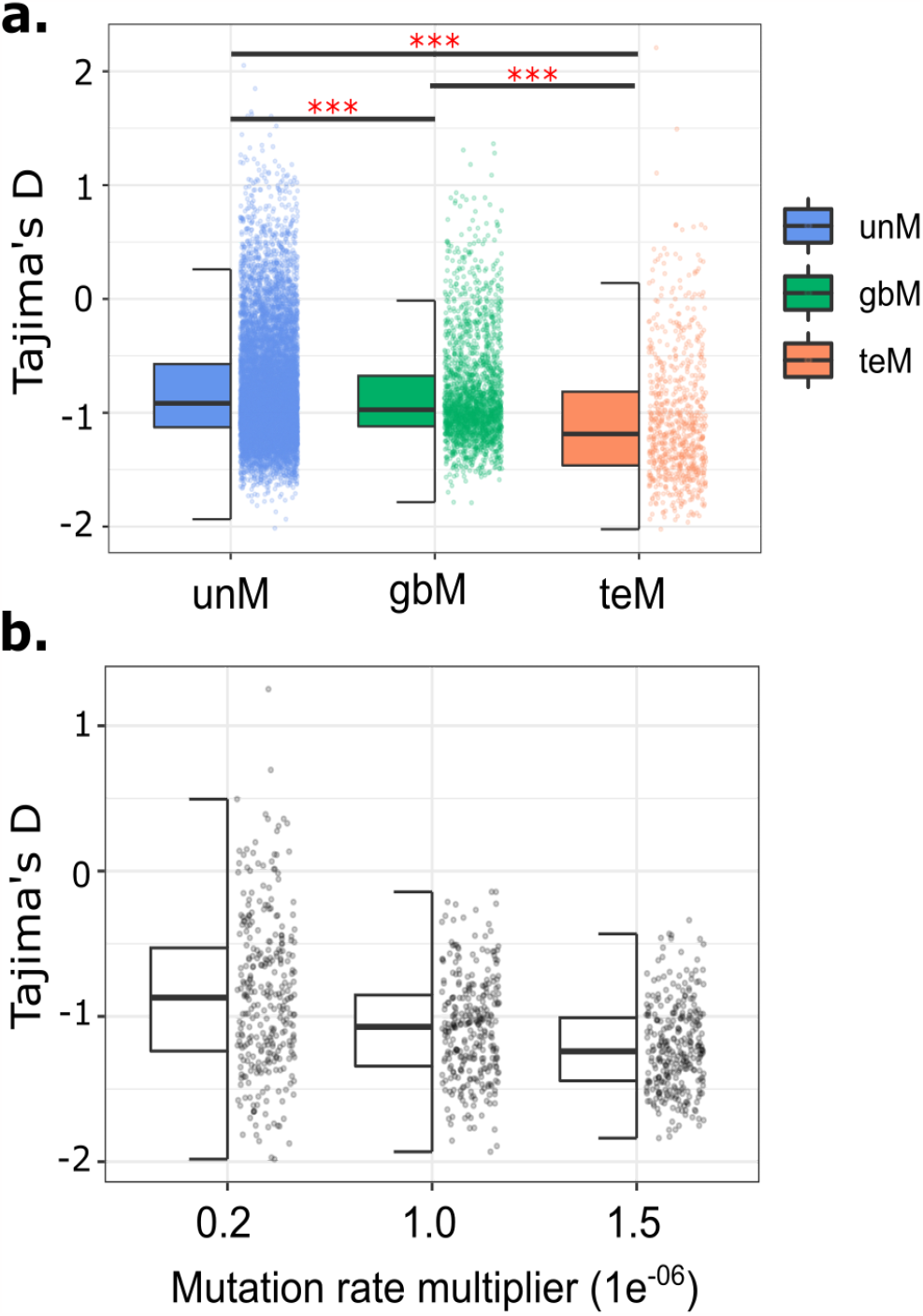
The site-frequency spectrum, Tajima’s D, for different classes of genie methylation. **a)** Distribution of population genetic test statistic, Tajima’s D, in unM, gbM, and teM genes. ’***’ denotes statistically significant difference in Tajima’s D in pairwise comparision following a one-way ANOVA test at p-value < 0.001 **b)** Simulation of Tajima’s D from 3000 bp regions using a population of 1000 individuals and 10000 generations replicated 300 times for different mutation rates (0.2, 1.0 and 1.5. e.g., 1.5 = 150% increase in mutation rates).

To better interpret observed population genetic statistics, we conducted evolutionary simulations modeling a 3000 bp region under different mutation rates (0.2x, 1x, 1.5x) deleterious mutation frequency (0%, 10%, 20%, and 50%), and strength of selection (0 to -0.5). As anticipated by existing theory, more negative Tajima’s D values were observed as an effect of higher mutation rates (1.5 × 10^−06^ bp^-1^ generation^-1^) or a higher proportion of deleterious mutations (50%) (One-way ANOVA, f=60.82, p-value < 0.001; f=33.71, p-value < 0.001, respectively) (**Figure 2b, S4**). The more negative Tajima’s D observed in gbM genes compared to unM genes is consistent with the effects of purifying selection. In contrast, the considerably high magnitude of negative Tajima’s D in teM may be due to their increased mutation rate and cannot be explained by teM genes having more sites under selection. Furthermore, differences in the distribution of fitness effects could, in principle, also cause more negative Tajima’s D and a higher number of polymorphisms (**Figure S5**) (Charlesworth and Jensen 2022). For example, under a specific set of conditions, Tajima’s D can be more negative in genes where selection is at intermediate strength compared to genes where selection is extremely strong (all selected mutations are nearly lethal). However, in comparisons of gene duplicates distinguished by origin (WGD vs tandem, etc) rather than by epigenome states, confirm that more negative Tajima’s D is associated with stronger purifying selection (**Figure S6, S7**) Therefore, both differences in selection and mutation rate (increased mutation rates in teM genes) jointly explain the relative differences in signatures of selection, polymorphisms, and Tajima’s D among gbM, unM, and teM genes. These findings are, therefore, consistent with a significant impact of polymorphism variability via mutation rate and selection variability on sequence evolution among genes defined according to epigenome state.

### Epigenome-mediated mutation rate differences in paralogs with divergent methylation

DNA methylation plays an important role in the expression and evolution of duplicate genes (Adams 2007; He et al. 2022). Previous work has shown that paralogs with divergent methylation show more divergence in expression patterns than paralogs with similar methylation states (Raju et al. 2022). We classified *A. thaliana* genes based on their types of duplication into whole-genome duplicates, tandem, proximal, translocated (also known as transposed), and dispersed duplicates using *DupGen_finder-unique (Qiao et al. 2019)*. Whole-genome duplicates (WGDs) had lower polymorphism (One-way ANOVA, f(6)=106.8, p-value < 0.001; **Figure S6**) compared to the four different types of single-gene duplicates (SGDs). Within SGDs, non-syntenic SGDs such as translocated and dispersed duplicates had lower polymorphism in the natural accessions compared to local SGDs: tandem and proximal (Tukey’s test, adj p-value < 0.001; **Figure S6**). WGDs also showed more negative Tajima’s D values than local SGDs: tandem and proximal duplicates (Tukey’s test, adj p-value < 0.001, **Figure S6**). The lower rates of polymorphism in WGDs combined with a more negative Tajima’s D values indicate stronger purifying selection on WGDs consistent with their lower Dn/Ds ratio (**Figure S7**) compared to SGDs, particularly tandem and proximal duplicates.

To further explore how DNA methylation divergence influences the evolutionary fates of duplicate paralogs, we compared duplicate pairs based on their genic methylation states (**Table S2**). Previous work has shown that expression divergence between duplicate paralogs is much lower in pairs with similar methylation profiles than those with genic methylation divergence (Raju et al. 2022). Thus, duplicate pairs that retained the same methylation profile were excluded, and 1010 duplicate gene pairs (677 unM-gbM, 298 unM-teM, and 35 gbM-teM pairs) that showed divergence in genic methylation states were included for further analysis. In unM-teM and gbM-teM pairs, the teM paralogs showed a significantly higher percentage of polymorphism in natural populations compared to the unM and gbM paralogs (T.test, p-value < 0.001; **Figure 3a**). In contrast, we did not find any significant difference in the percentage of polymorphism between unM and gbM paralogs in unM-gbM pairs (T-test, p-value = 0.198). We further explored if the population genetic test metric Tajima’s D also shows corresponding differences between duplicate pairs. TeM paralogs in unM-teM pairs showed significantly more negative Tajima’s D values compared to its unM paralog (T-test, p-value < 0.001; **Figure 3b**), consistent with elevated mutation rate or differences in the DFE between these genes. Differences were not observed in the other pairs, consistent with interactive effects of selection and mutation rate differences. The directionality of changes in methylation states cannot be typically ascertained for most duplicate pairs. However, in the case of translocated duplicates, the parental (ancestral) copy is found in synteny with other species (Wang et al. 2013). Thus we can discern the direction of methylation change by assuming the methylation at the parental locus as the original state. Translocated paralogs that gained teM methylation showed higher polymorphism in *A. thailana* accessions compared to their parental unM and gbM paralogs (**Figure 4**, ttest, p-value = 0.0152 and pvalue < 0.001, respectively). Conversely, when teM parental paralogs switched to unM translocated paralogs, they showed lower polymorphism (ttest, p-value = 0.0027). Collectively, these results indicate that similar to the epigenome-mediated polymorphism biases observed at whole genome scales, duplicate paralogs that diverge in their epigenomic states also show comparable differences in polymorphism.

**Figure 3:**
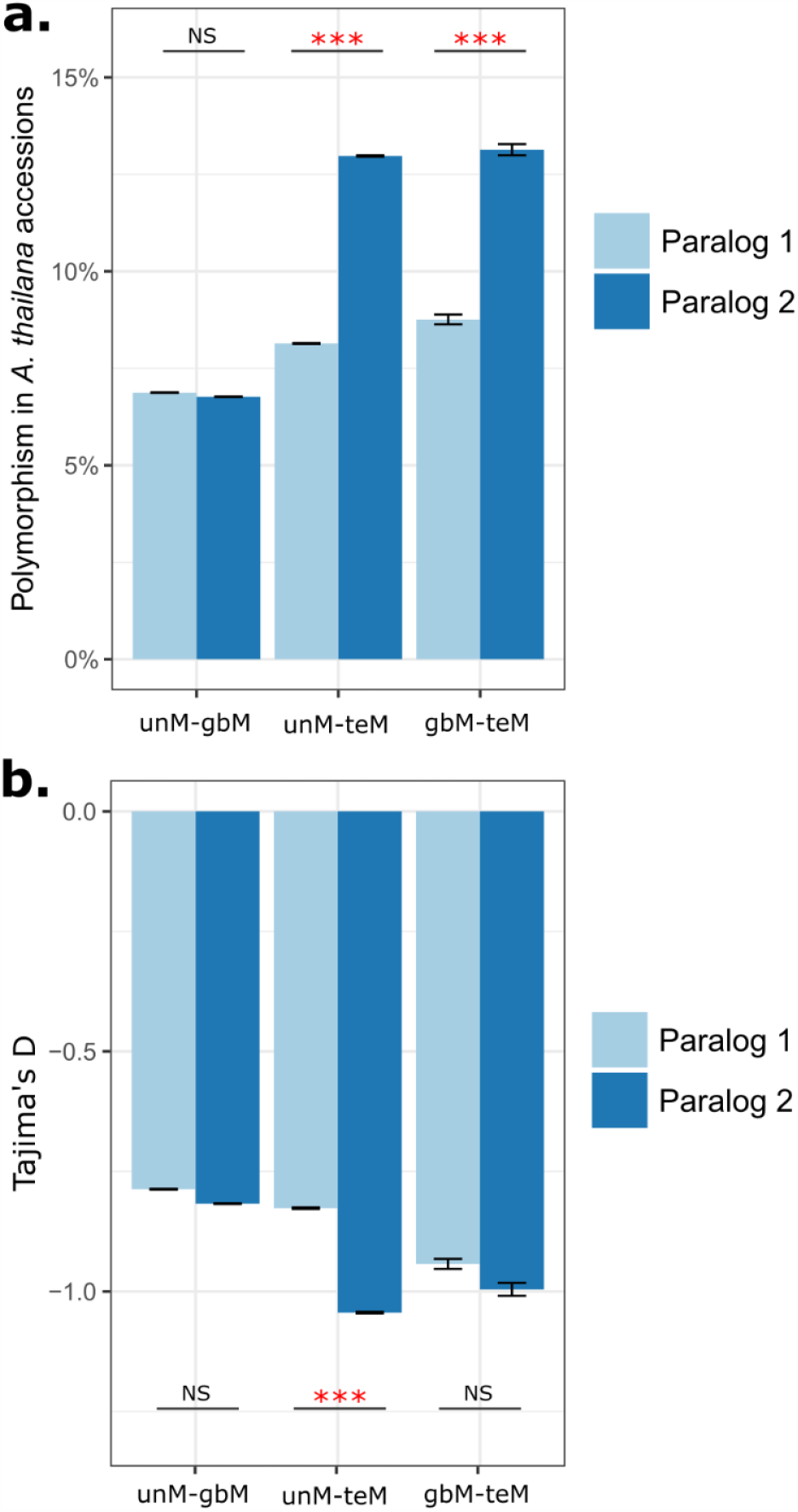
Epigenome mediated polymorphism differences in duplicate paralogs. **a)** Percent polymorphism in duplicate gene pairs with methylation divergence. For example, a duplicate pair with a unM paralog and a gbM paralog are classified under unM-gbM (n = 677 pairs). Similarly for unM-teM (n = 298 pairs) and gbM-teM (n = 35 pairs). Hest was performed to test differences in polymorphism between paralogs. Error bars represent standard error of mean and ’***’ represents a statistical significance at p-value < 0.001. **b)** Mean differences in the population genetic test statistic, Tajima’s D in duplicate gene pairs with methylation divergence. Hest was performed to test differences in Tajima’s D between paralogs. Error bars represent standard error of mean and ’***’ represents statistical significance at p-value < 0.001. NS denotes ’Not statistically significant’.

**Figure 4:**
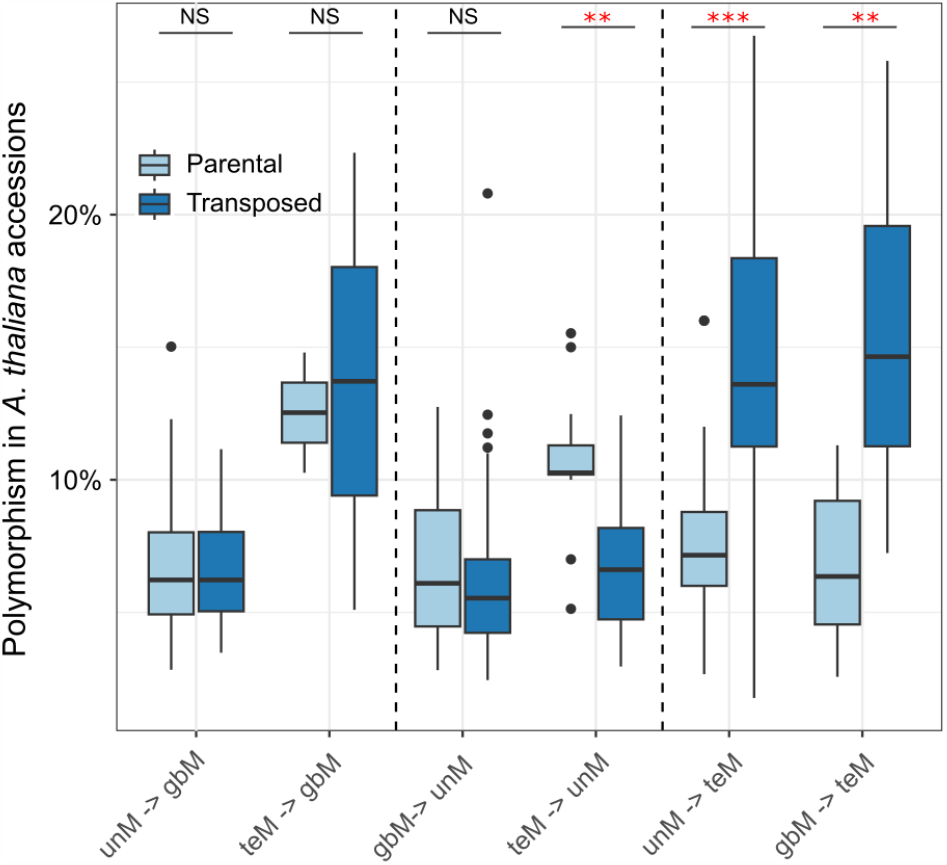
Polymorphism rates correspond well with the direction of methylation chages in transposed duplicates. Percent polymorphism in *A. thaliana* accessions in ancestral and translocated duplicate pairs ordered based on their methylation state switching. For example in unM -> gbM, the ancestral paralog is unmethylated while the translocated paralog has gained gene-body methylation. At-test was performed to test for differences in polymorphism rates between ancestral and translocated duplicate pair. ’***’ represents statistical significance at p-value < 0.0001, and ’**’ for p-value < 0.05.

### TeM paralogs show depletion of H3K4me1 and targeted DNA repair protein PDS5C

To consider the potential role of mutation rate variation in more detail, we examined divergence in DNA repair and H3K4me1 in gene duplicates. Previous work has shown that PDS5C, a cohesion co-factor protein involved in homology-directed repair (HDR), targets H3K4me1 associated with gene bodies of transcriptionally active and evolutionarily conserved genes (Quiroz et al.). Because the Tudor domain responsible for H3K4me1 binding in PDS5C is also found in the mismatch repair protein MSH6, the distribution of PDS5C may serve as a proxy for repair activity generally. Indeed, knockout lines of MSH6 and its partner MSH2 show elevated mutation rates in H3K4me1-marked regions (Belfield et al. 2018; Quiroz et al. 2022). Here we tested whether duplicate paralogs with different epigenomic states show differences in their association with the H3K4me1 histone mark and DNA repair protein PDS5C. TeM-containing paralogs have significant depletion of H3K4me1 compared to their unM and gbM paralogs. In contrast, gbM-containing paralogs have significant enrichment of H3K4me1 compared to unM and teM paralogs (t-test, p-value < 0.001) (**Figure 5a**). These differences between paralogs further confirm the strong association of H3K4me1 with gene-body methylation (Fig S1). Similarly, we find that PDS5C targeting is depleted in teM paralogs while PDS5C is increased in gbM and unM paralogs (t-test, p-value < 0.001) (**Figure 5b**). These results provide a potential mechanism for the divergence of duplicate paralogs, where divergent genic methylation is associated with histone marks of paralogs that influence preferential targeting of DNA repair proteins consequently leading to divergence in rates of sequence polymorphism (mutation) and different evolutionary fates of duplicate genes.

**Figure 5:**
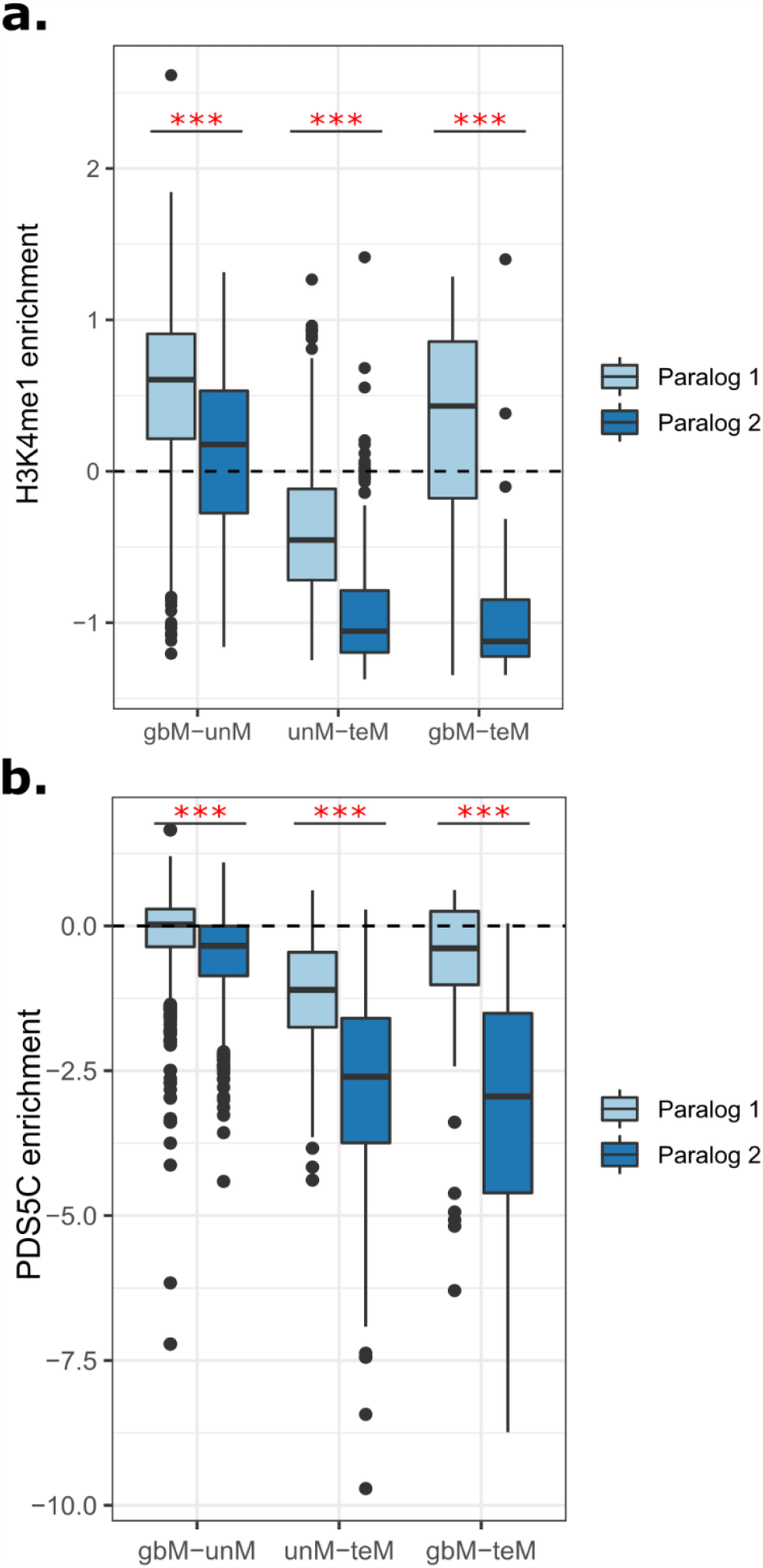
Paralogs show differential enrichments of the histone mark H3k4me1 and DNA repair protein PDSSC. **A)** Difference in H3k4mel enrichment in duplicate paralogs with methylation divergence. T-test was performed to test differences in H3K4mel enrichments between paralogs. *** represents a statistical significance at p-value < 0.001. **B)** Enrichment of PDSSC, a cohesion cofactor gene involved in homology-directed repair (HDR). T-test was performed to test differences in PDSSC enrichments between paralogs. *** represents a statistical significance at p-value < 0.001

## Discussion

We investigated whether the evolution of gene expression patterns and epigenomic states correspond with changes in the levels of polymorphism between duplicate paralogs in *A. thaliana*. First, we compared *A. thaliana* genes with distinct epigenomic states previously shown to correspond to different expression patterns: unmethylated (unM), gene-body methylated (gbM), and transposon-like methylated genes (teM) and showed that CG-only methylated gbM genes tend to accumulate fewer polymorphism than unM genes. However, the relationship between accumulation of polymorphism and DNA methylation is complex since teM genes with CG and non-CG methylation show higher polymorphism than unM and gbM genes.

To more precisely compare rates of sequence polymorphism, we focused on duplicate paralogs with divergent methylation states. Comparing duplicated genes following different trajectories allows an internally controlled test of the link between epigenomic state, expression level, and potential variation in polymorphism rate/divergence. (**Figure 6**). Previous work has shown that DNA methylation states are associated with differences in expression specificity, measured as the breadth or narrowness of gene expression patterns across multiple tissues/conditions (Keller and Yi 2014; Wang et al. 2014; Kim et al. 2015; Raju et al. 2022). Our results show that paralogs with silenced epigenomic states (teM) tend to have higher polymorphisms across the natural population compared to their active paralogs (unM and gbM). These observations of polymorphism rate changes apparent in both genome-wide and duplicate paralog comparisons provide insights into the evolutionary processes reflecting the intertwined relationships between gene expression, epigenomic states, and mutation rates.

**Figure 6:**
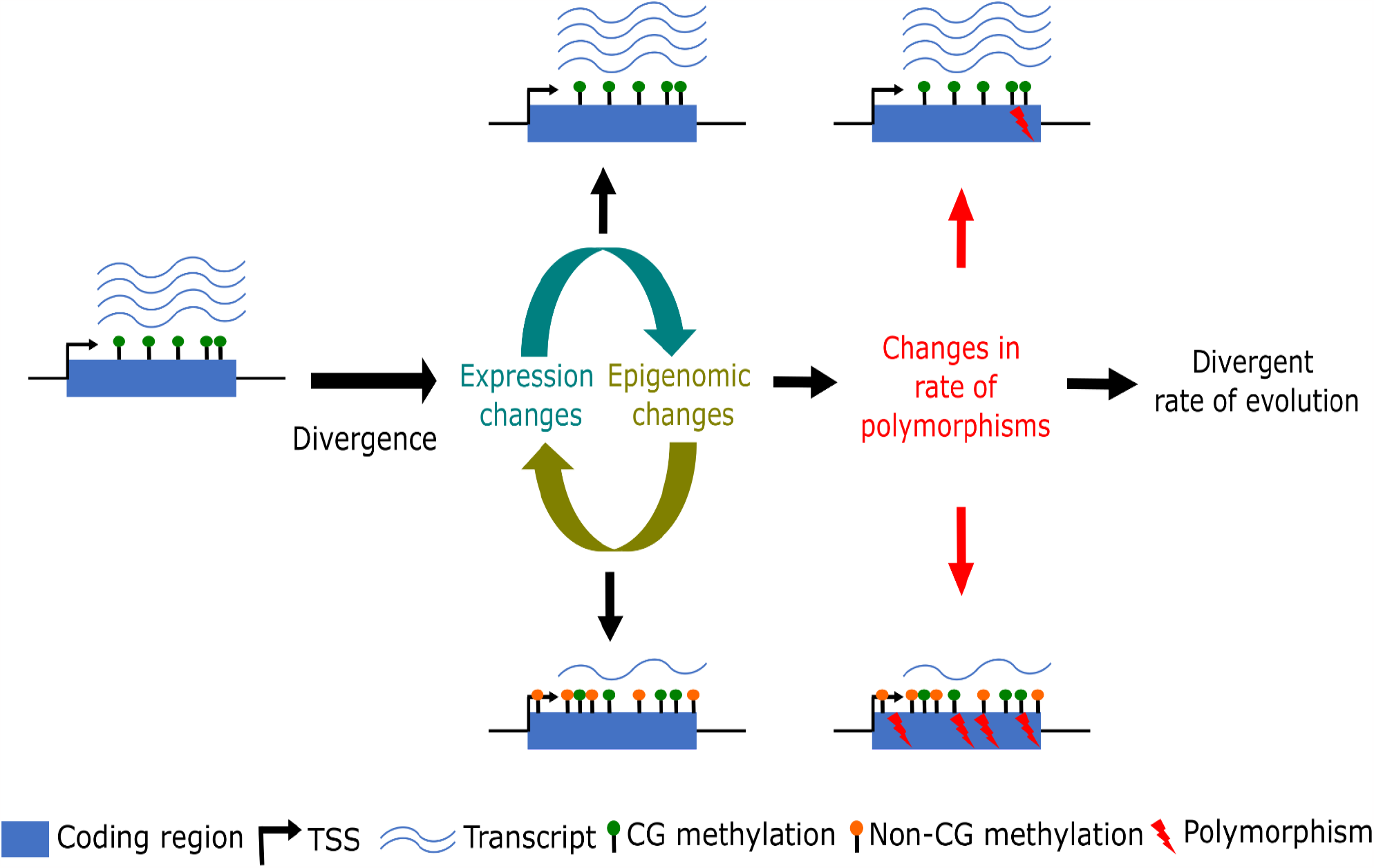
Gene expression and epigenomic divergence leads to different rates of polymorphisms in duplicate paralogs. Schematic representation showing gene expression and epigenomic changes in duplicate paralogs leading to changes in polymorphism rates and subsequently divergent evolutionary fates of duplicate genes.

Whether these sequence polymorphisms are independent of gene activity or not is important in predicting the evolutionary fate of duplicate paralogs and the maintenance of genetic redundancies. Nowak et al. 1997 developed genetic models to examine selection pressures on redundant genes (Nowak et al. 1997). Based on their models, redundant genes remain evolutionarily stable when

1) both paralogs perform a function with equal efficacy and have equal mutation rates, 2) the paralog that performs a function with lower efficacy has lower mutation rates, and 3) one of the paralogs gain a new function (neofunctionalization). Divergence in paralog expression specificity is known to be a major factor in the retention and maintenance of paralogs via sub- and neo-functionalization (Duarte et al. 2006), and gene copies have been shown to change expression immediately following duplication (Comai et al. 2000; Shaked et al. 2001). This is particularly true of SGDs, especially those that are not copied in their local chromatin environment and showed DNA methylation and expression divergence (Ganko et al. 2007; Raju et al. 2022).

Our results show these gene expression and methylation divergence across the genome and between paralogs are associated with differences in observed polymorphism. Moreover, polymorphism rates are higher in the epigenomically silenced teM paralogs, associated with low and tissue-specific expression, compared to the active gbM paralog, associated with medium to high expression and a much broader expression pattern across different tissue/conditions (Zhang et al. 2006; Zilberman et al. 2007). Both these observations limit the frequency in which models 1 and 2 likely explain the evolutionary stability of redundant paralogs following duplication within this system (Nowak et al. 1997). Redundancy in gene copies can be maintained if one paralog gains a new function, as outlined in model 3. Based on our results, sub- and neo-functionalization are a more likely path for the retention of gene duplicates rather than the equilibria explained by models 1 and 2. While our results show polymorphism rates to be qualitatively associated with epigenomically active (gbM) and repressed (teM) states, more modeling and simulation studies are needed to quantify how different epigenomic and expression states influence observed differences in polymorphism between paralogs.

The existing theory also anticipates that the elevated polymorphism (if via elevated mutation rates) observed in silenced genes could help facilitate the origins of novel functions. Deleterious emergent functions could constrain new mutations in a paralog that alter its function. However, if hypermutating paralogs are those that are also silenced epigenomically, the negative consequences of deleterious mutations could be reduced, allowing for the exploration of a broader functional landscape, providing time for novel functions, potentially even across fitness valleys, to emerge (Krakauer and Nowak 1999; Rodin and Riggs 2003; Rodin et al. 2005). Epigenomic silencing is usually found in a tissue-, condition- and/or development-specific manner favoring functional divergence between duplicates (Adams et al. 2003). It is conceivable, therefore, that the relationship between mutation rate and expression across the genome and following duplication allows transiently silent but reactivable genes to neofunctionalize and later transition to a transcription-activated state and conferring some fitness advantage benefitting from a reduced mutation rate (Rodin and Riggs 2003; Rodin et al. 2005). Still, while our findings support the potential for these dynamics, future work is needed to test these more quantitatively and characterize specific test cases, for example, testing if there is any bias towards neutral or positive mutations in epigenomically, silenced genes.

Housekeeping genes with broad expression patterns are known to be enriched in CG-only methylated gbM genes. This essentiality could explain the selection of mechanisms facilitating lower observed polymorphism in gbM genes. GbM genes are expressed at much higher levels than teM genes, and actively transcribed genes are protected by DNA repair machinery during replication and transcription, leading to lower mutation rates (Gonzalez-Perez et al. 2019). However, the epigenomic environment around the gene influences the recruitment of DNA repair machinery to actively transcribed genes (Frigola et al. 2018). For example, in the human genome, the histone mark, Histone H3K36me3, marks gene bodies and recruits the mismatch-recognition protein MutSα (Huang et al. 2018). In plants, gene bodies are marked by other histone modifications, such as H3K4me1, previously shown to be associated with lower mutation rates (Monroe et al., 2022; Quiroz et al. 2022). H3K4me1 marks are enriched in gbM genes/paralogs and show depletion in teM genes/paralogs. Other factors, such as chromatin organization and replication timing, can also contribute to mutation rate differences (Stamatoyannopoulos et al. 2009; Schuster-Böckler and Lehner 2012), though the established theory has demonstrated that these are more likely to be the result of netural processes rather than the kind of selective pressures capable of shaping links between epigenomic features and mutation rate (Lynch 2010; Martincorena and Luscombe 2013; Lynch et al. 2016). A plausible mechanistic model is that the segmentation of duplicate paralogs into epigenomically active and silent states captures divergence in the activity of DNA repair mechanisms resulting in mutation rate variation in relation to epigenomic states, with resulting mutations then being subject to selection. Patterns of sequence variation in these gene pairs may thus reflect the joint action of mutation rate and selection variation.

In summary, we find that observed polymorphism in *A. thaliana* genes is not independent of expression-associated epigenomic states. Considering these relationships, we see implications for the evolutionary fate of gene duplicates, suggesting a role in promoting the origin of novel gene function. Still, future work is needed to test emergent hypotheses. Elucidating the precise mechanisms and evolutionary origins of these patterns will contribute to a deeper understanding of the reciprocal relationship, potentially influenced by evolved mutational processes, between evolutionary and molecular biology.

## Acknowledgments

This work was supported by the University of California Startup Funds and Department of Plant Sciences (JGM, ML, DK), Michigan State University Startup Funds (CN), and the National Science Foundation (grant IOS-2029959 to CN).

## Author Contributions

SKKR, GM, and CN designed the study and analysis. SKKR and ML performed data analysis. DJK contributed to the interpretation of the results. SKKR, GM, DJK, and CN wrote and edited the manuscript. All authors read and approved the final manuscript.

## Data availability

All data and codes to perform data analysis and generate plots for figures are available on GitHub (https://github.com/Kenchanmane-Raju/Ath_Paralog_Divergence)

## Competing interests

The authors declare that they have no competing interests.

**Figure S1:**
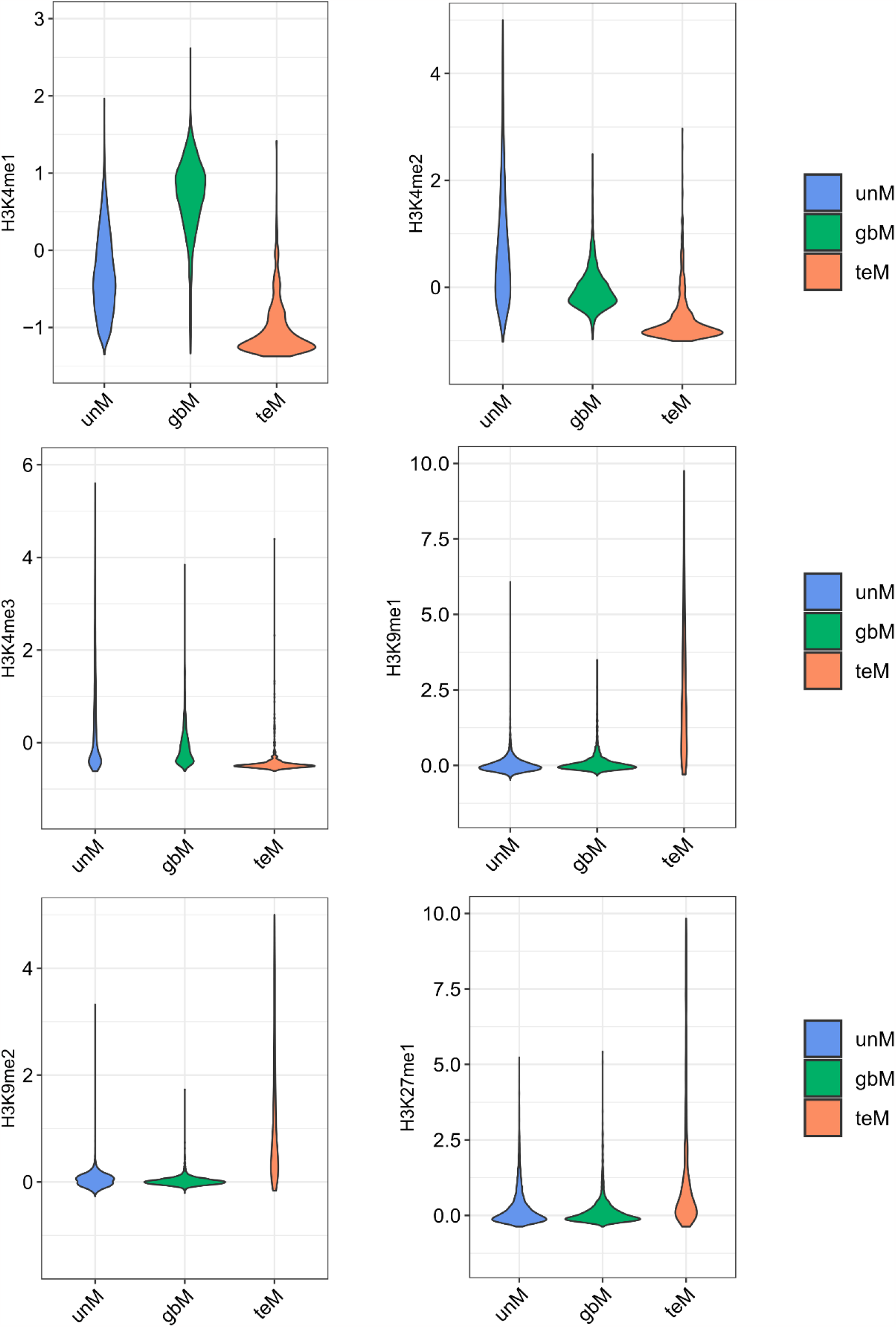
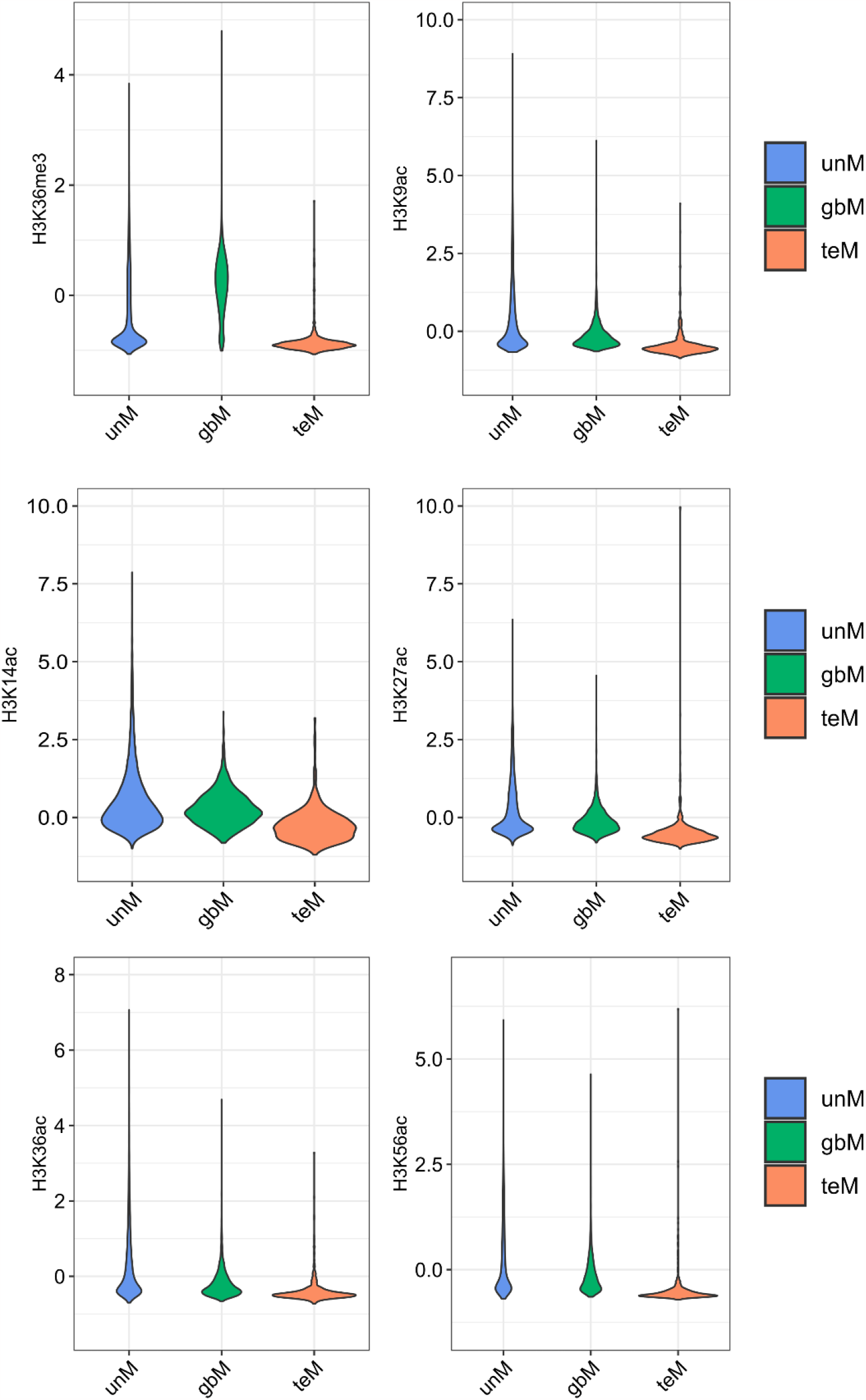
Histone modification marks in different genie mthylation classes (unM, gbM, and teM). Each histone modifaction was scaled and averaged across the genie region.

**Figure S2:**
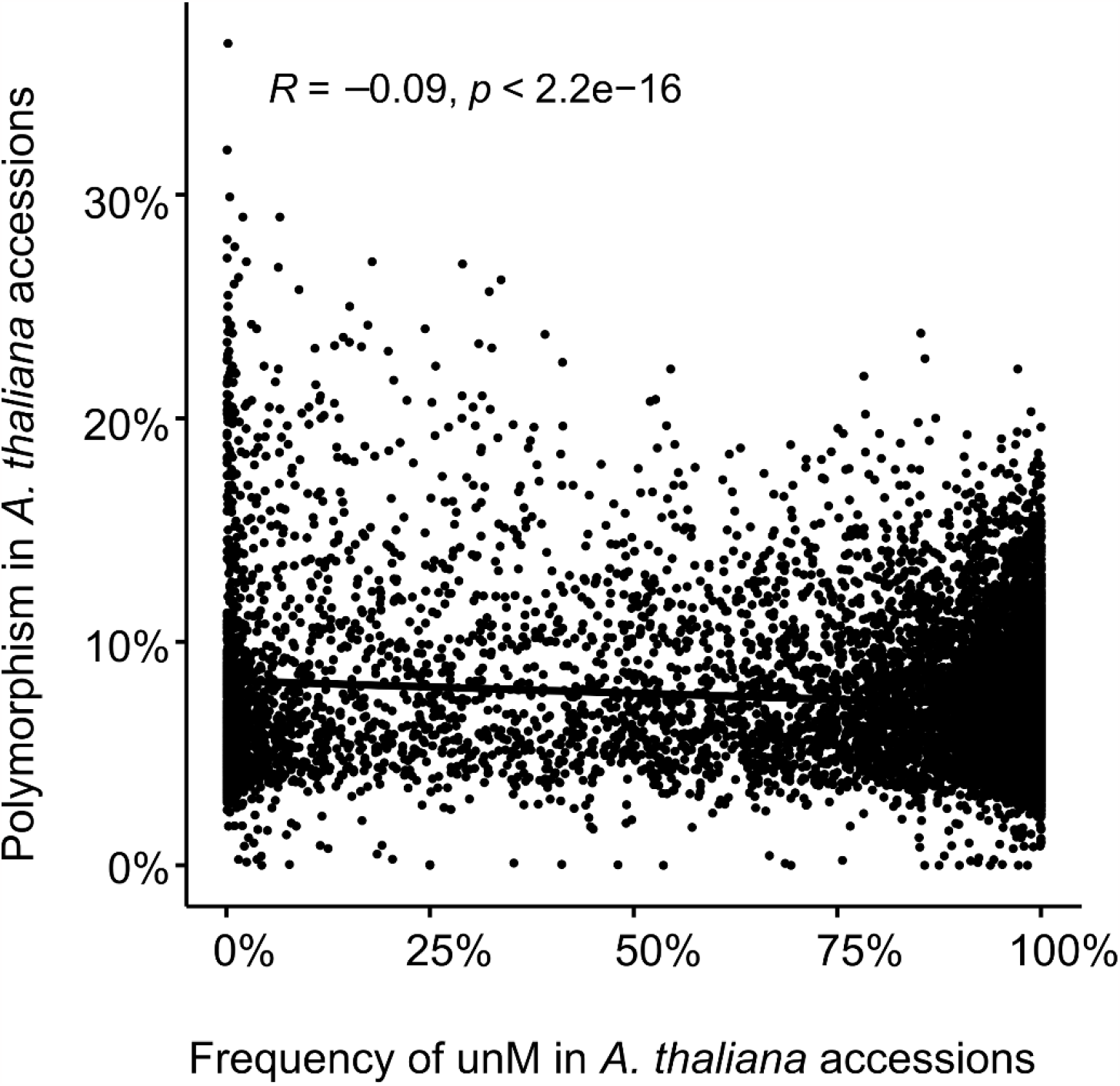
Spearman’s rank order correlation between frequency of unM epiallele and single-nucleotide polymorphism in the natural accessions of *A. thaliana*. Negative Spearman’s rho (r_s_) is indicated.

**Figure S3:**
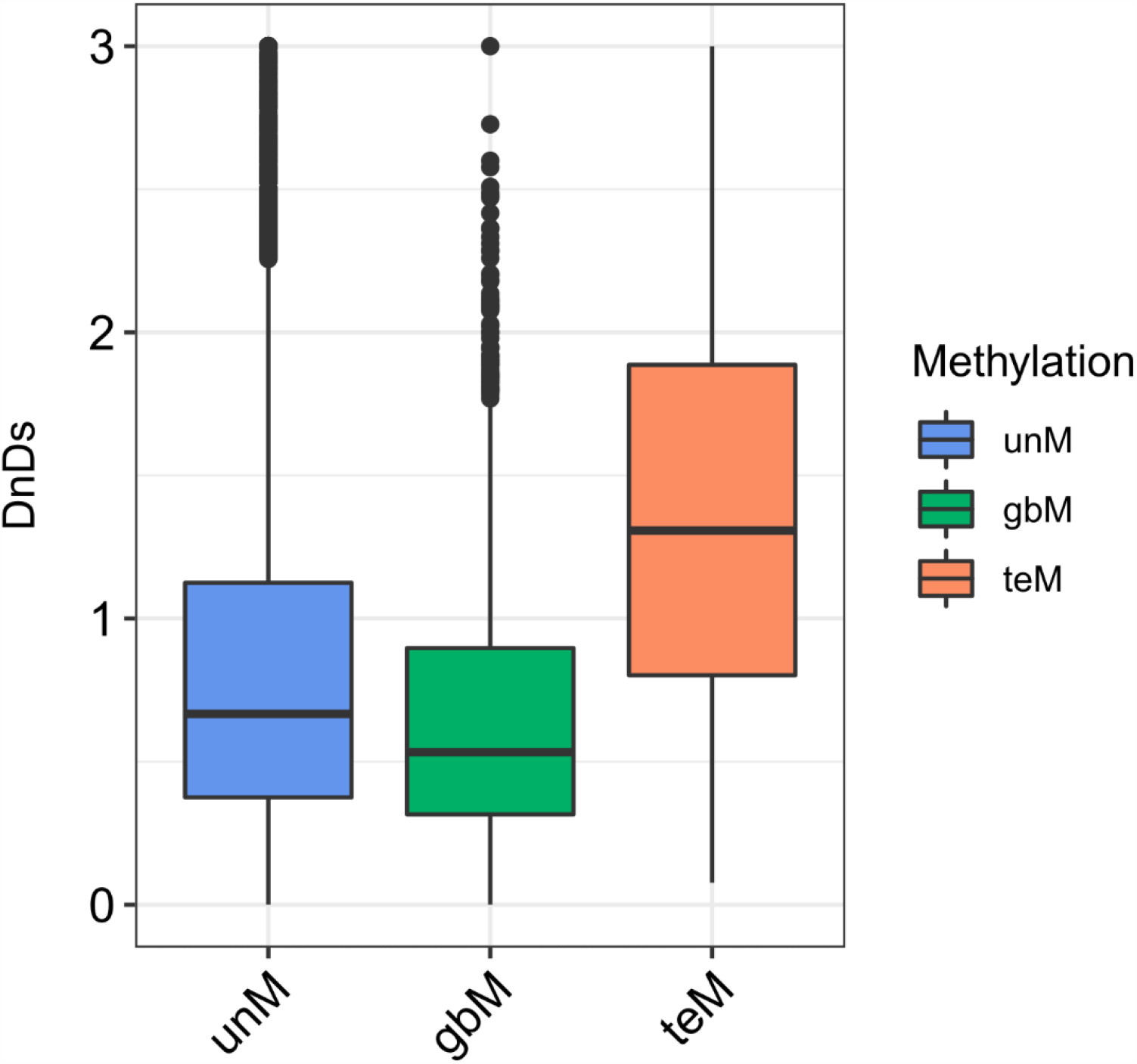
Signatures of selection among the three genie methylation classification (unM, gbM, and teM). Dn and Ds ratios were calculated as non-synonymous (Dn) and synonymous (Ds) substitution divergence from *Arabidopsis lyrata*.

**Figure S4:**
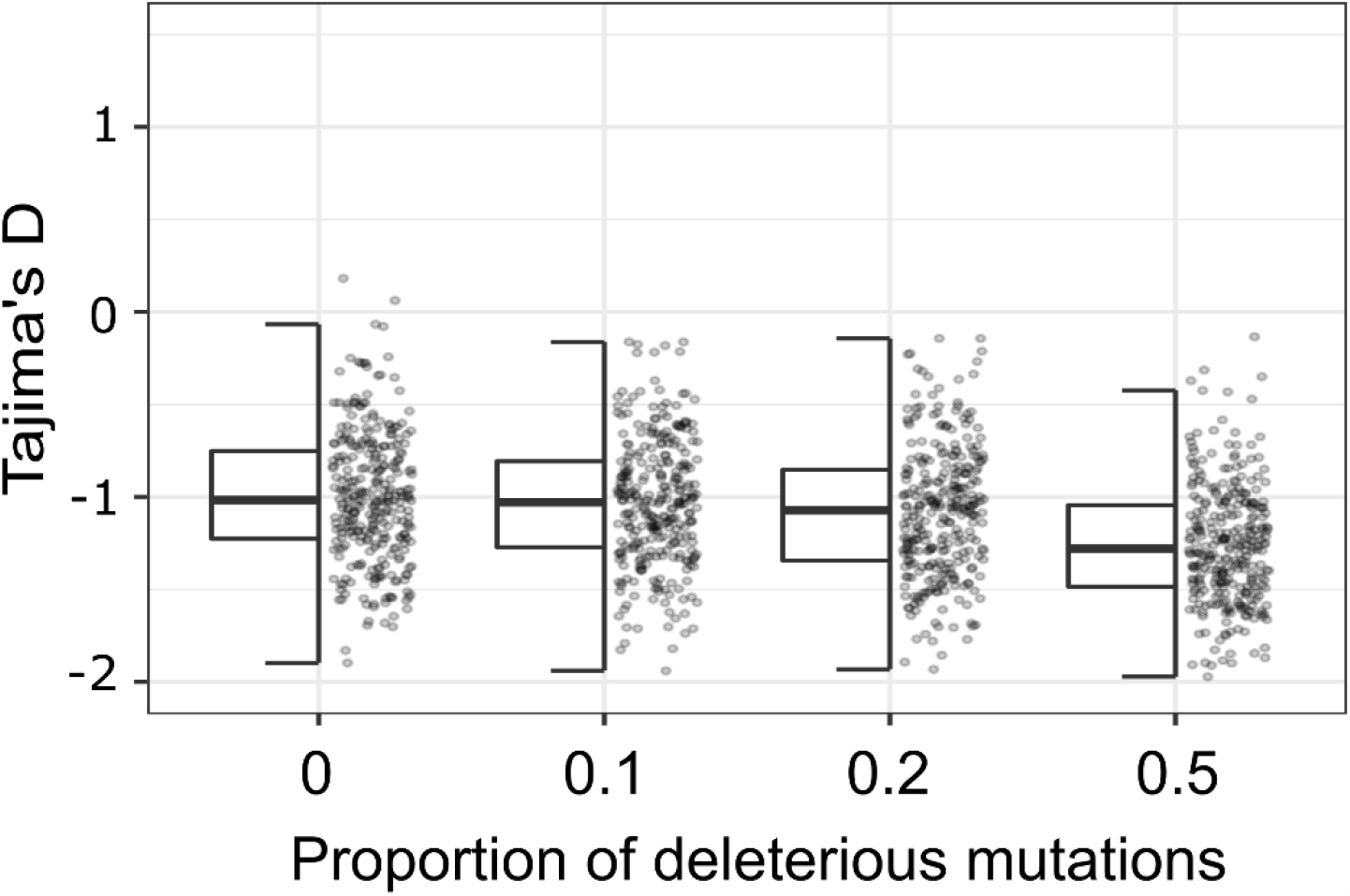
Simulation of Tajima’s D from 3000 bp regions using a population of 1000 individuals and 10000 generations replicated 300 times for different proportions of deleterious mutations (0, 0.1, 0.2, and 0.5 (50% deleterious mutations)).

**Figure S5:**
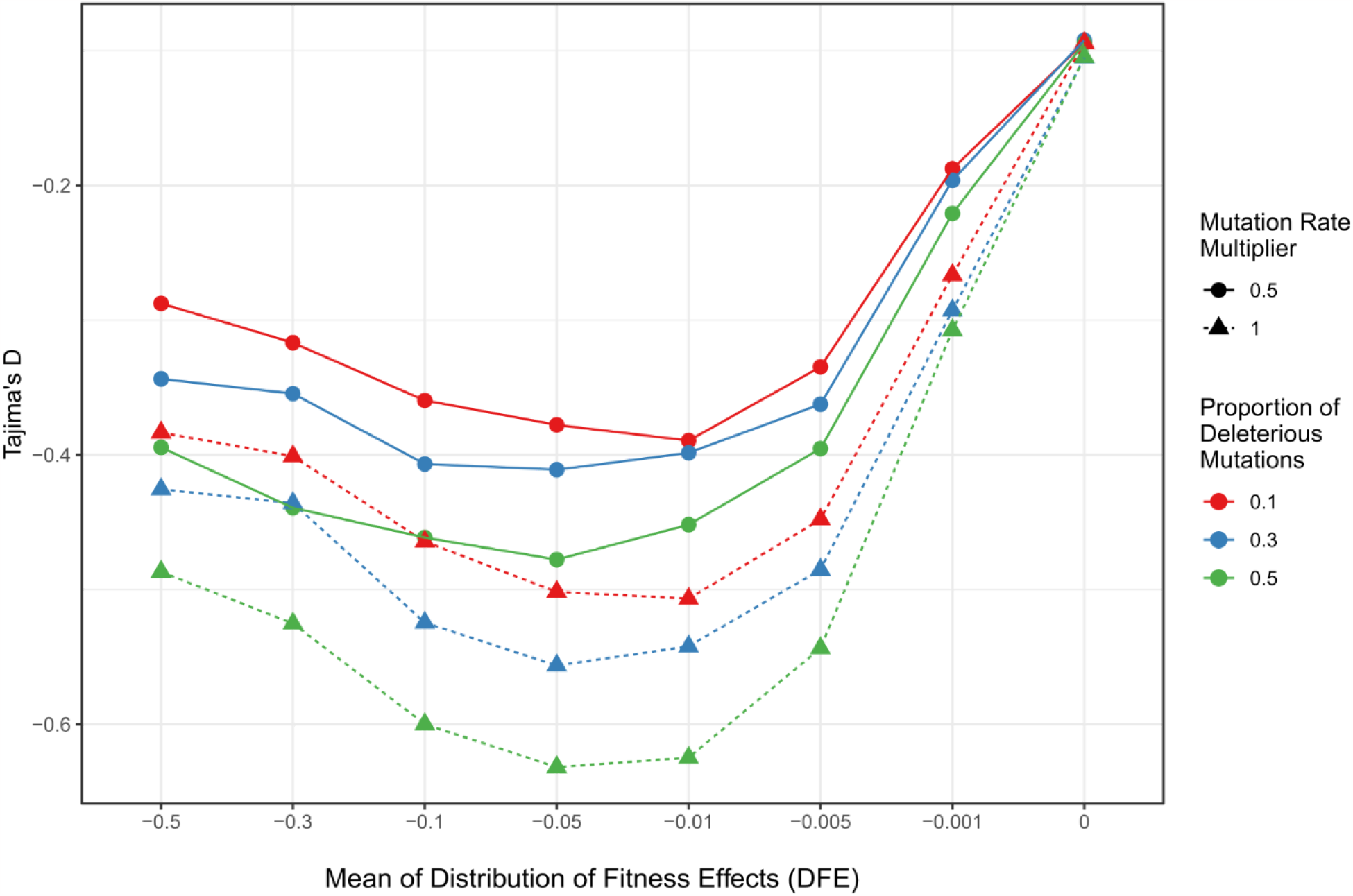
Effect of mean value of distribution *effect* (DFE) on Tajima’s D. Each point represents the average Tajima’s D value for a unique parameter combination simulated from a 3,000 bp sequence in a population of 1,000 individuals after 10,000 generations in 300 replicates.

**Figure S6:**
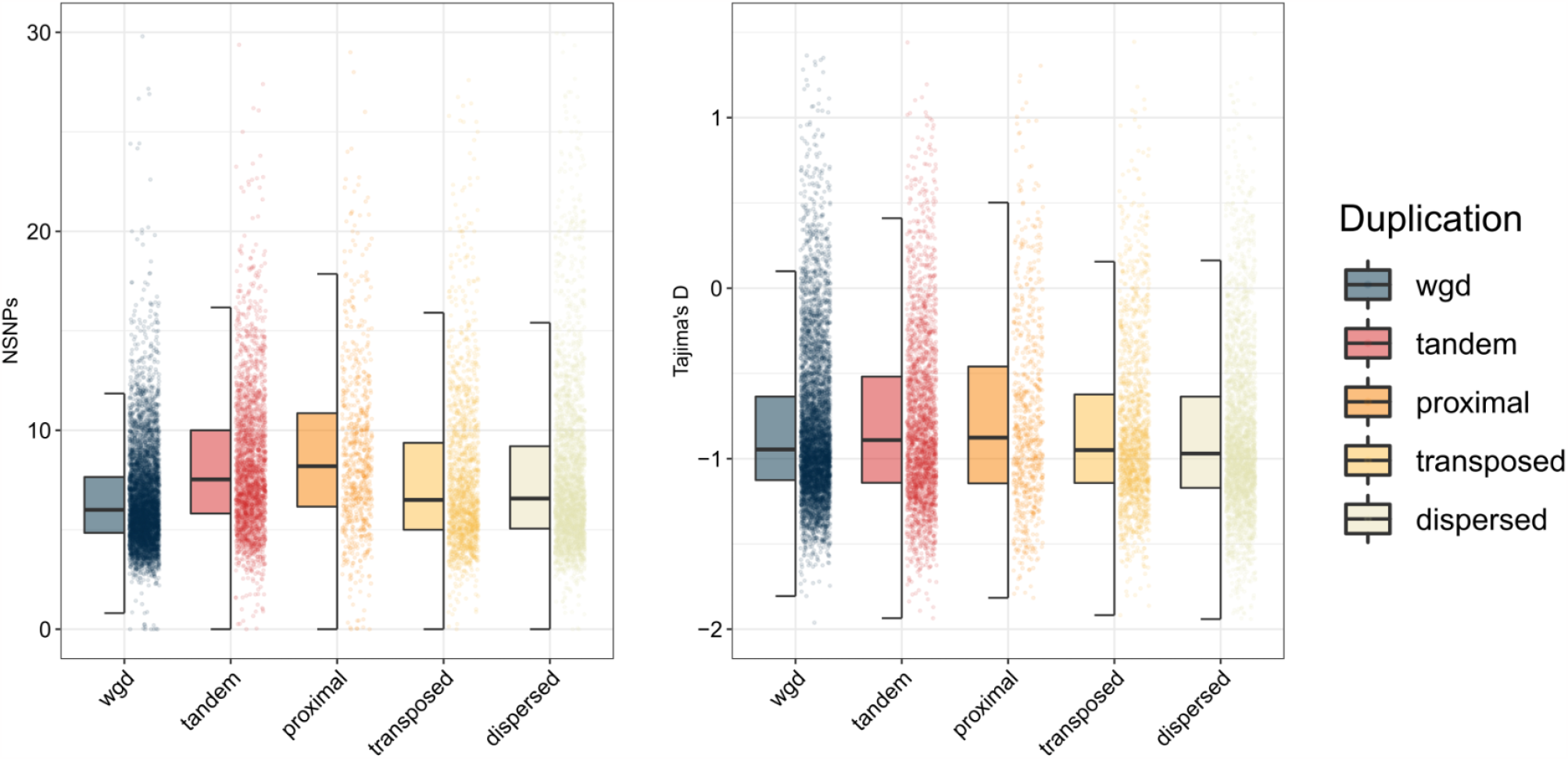
Rate of polymorphism and Tajima’s D differences between different types of gene duplicates. Percentage of nucleotides with single nucleotide polymorphisms in *A. thaliana* accessions measured in 100 bp windows averaged for each gene belonging to different types of gene duplicates. Population genetic statistic, Tajima’s D values for different types of duplicates. WGD - whole genome duplicates. Tandem, proximal, translocated, and dispersed are different types of single-gene duplicates.

**Figure S7:**
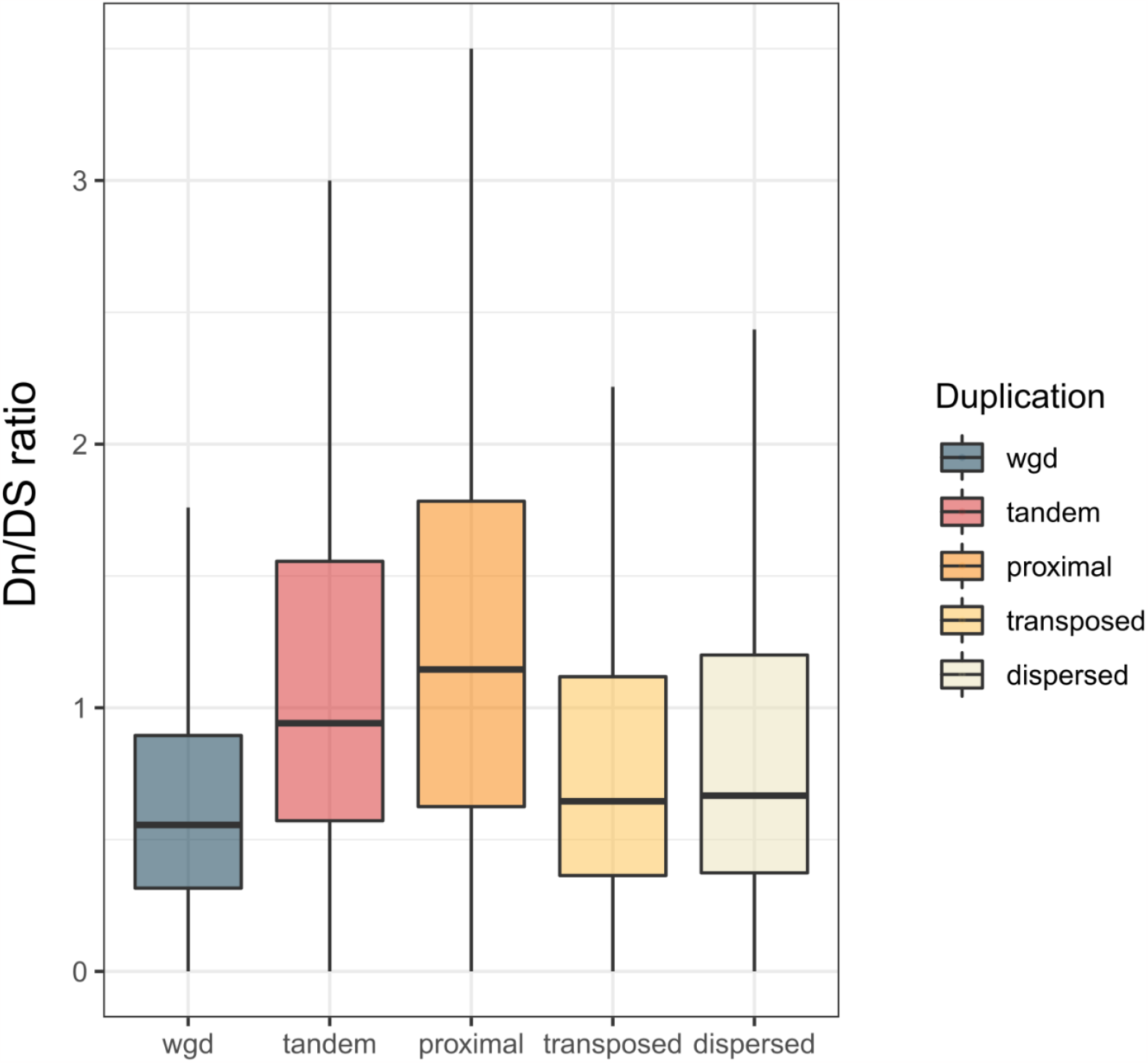
Dn/Ds ratios of different types of gene duplicates. Ratio of signatures of selection for each A. thaliana gene from different modes of gene duplication, using their synonymous (Ds) and non-synonymous (Dn) substitution divergence from Arabidopsis lyrata.

